# Temperature variation as a driver of *Wolbachia* release efficacy: implications for dengue control in warming climates

**DOI:** 10.64898/2026.07.26.740614

**Authors:** Juanita Prada-Mora, Santiago A. Villamil-Chacón, Mauricio Santos-Vega

## Abstract

**Background:** *Wolbachia*-based control methods reduce dengue transmission by suppressing *Aedes aegypti* populations or blocking viral replication, yet their effectiveness across the climatic conditions of endemic regions remains poorly understood. Seasonal and interannual temperature changes shape mosquito dynamics, but how *Wolbachia* releases perform during anomalous events such as El Niño or heatwaves is largely unknown—a gap that limits our ability to optimize release strategies and predict intervention success across different climates.

**Methodology/Principal Findings:** To address this gap, we built a mathematical modeling framework that explicitly incorporates temperature-dependent *Wolbachia* parameters together with the seasonal and interannual climate variability characteristic of dengue-endemic regions, coupling a detailed *Wolbachia* dynamics model with a susceptible–infectious–recovered (SIR) epidemiological model to evaluate *Wolbachia* establishment, persistence, and stability under different thermal regimes and trace their downstream impact on disease spread. Higher temperatures eroded both *Wolbachia* establishment and long-term persistence, sharply narrowing the range of effective release strategies as conditions approached 30 °C. Seasonality added a further layer of complexity: the timing of thermal stress relative to *Wolbachia* frequency, not merely its magnitude, determined whether population replacement succeeded. Interannual shifts, progressive warming, widening seasonal swings, and displaced thermal peaks, each eroded *Wolbachia* prevalence and stability, with effects that compounded over successive years. Dengue transmission tracked these dynamics closely, with warmer conditions producing larger, earlier outbreaks, and intervention success hinging on how release timing, frequency, and targeting were matched to local thermal conditions.

**Conclusions/Significance:** As extreme heat events become more frequent under climate change, release programs that ignore local thermal conditions risk falling short where dengue control is most needed. By elucidating the mechanistic interplay among temperature, *Wolbachia*, and dengue, our findings help refine *Wolbachia* release programs to suit different climatic conditions, thereby strengthening dengue control as climate variability intensifies.

**Author summary:** Dengue sickens hundreds of millions of people each year, and releasing mosquitoes carrying the naturally occurring bacterium *Wolbachia*, which blunts the virus’s spread, has become one of the most promising tools for fighting it. Yet one factor that decides whether these releases succeed or fail is routinely overlooked: temperature. We built a mathematical model linking *Wolbachia* biology and mosquito population dynamics to temperature, to ask a practical question: which release strategies work under real-world, changing climate conditions? Heat narrowed the margin for success. Higher temperatures shrank the range of effective interventions and slowed the replacement of wild mosquitoes with *Wolbachia*-carrying ones. Timing mattered just as much as intensity: releases launched before peak heat allowed *Wolbachia* to establish at higher levels, before thermal stress eroded its fitness benefits. And success was not permanent—growing year-to-year temperature swings could destabilize *Wolbachia* populations even after they had become established, threatening long-term disease control. The public-health stakes were stark: *Wolbachia* cut peak dengue cases by 54.6% at 25°C, but by only 10% at 30°C. For the communities most burdened by dengue, these results carry an urgent message. As climate change drives temperatures upward across endemic regions, *Wolbachia* programs designed without accounting for local thermal conditions risk underperforming precisely where they are needed most. Effective deployment requires climate-sensitive planning, adaptive release schedules, and continued investment in field-validated models that reflect the realities of a warming world.

## Introduction

*Aedes aegypti* is one of the most important vectors responsible for transmitting several arboviruses, including dengue, Zika, and Chikungunya [1]. In tropical and subtropical countries, the dengue virus (DENV) poses a significant public health issue, as dengue cases have risen notably in recent years, reaching a record high in the Americas in 2023 [2, 3]. In Colombia, DENV is hyperendemic across most regions, and all four serotypes have been reported throughout the country [4]. This co-circulation increases the risk of dengue hemorrhagic fever due to secondary infections. Strategies like vector control, community engagement, and surveillance aim to reduce transmission [4]. However, climate variations, such as changes in daily temperature, rainfall, and humidity, could lead to rapid increases in vector population density, and deficiencies in health services reduce the effectiveness of these measures, posing challenges for epidemiological analysis and control of dengue fever [5, 6].

Recently, in response to these challenges, *Wolbachia* interventions have emerged as effective, innovative strategies to reduce the dengue burden in regions such as South America, Asia, and Oceania [7–10]. *Wolbachia* is a Gram-negative, intracellular bacterium estimated to naturally infect about 60% of arthropod species [11]. It is maternally inherited and can alter host fitness and reproductive dynamics [12, 13], a property that can be harnessed for disease control. One of its most important properties is its ability to inhibit RNA virus replication, including dengue virus, by blocking pathogen dissemination within the host and thereby suppressing transmission [14–17]. When *Ae. aegypti* mosquitoes are transinfected with *Wolbachia*, they show substantially reduced vector competence for dengue virus, resulting in lower transmission rates and, in some cases, reduced mosquito population density [18, 19]. Together, these characteristics make *Wolbachia* a compelling tool for both suppressing and replacing wild vector populations in dengue-endemic settings.

The success of *Wolbachia*-based interventions relies on the bacterium’s capacity to spread and maintain itself within wild mosquito populations. *Wolbachia* gains a reproductive advantage via cytoplasmic incompatibility (CI) [20]. CI happens when uninfected eggs are fertilized by infected sperm, leading to non-viable offspring. Any male, infected or not, can fertilize infected eggs. This gives *Wolbachia*-infected females a reproductive advantage, boosting viable offspring and aiding the spread or replacement of wild populations, spreading the bacterium rapidly [20].

The first open-field trial of *Wolbachia* releases took place in Cairns, Australia, in 2011, in which Aedes mosquitoes and eggs were released into the environment, resulting in the successful integration of *Wolbachia* into the local mosquito population and the near-elimination of dengue transmission in treated areas [21]. Building on this, Indonesia implemented a suppression program using eggs and adult mosquitoes infected with the wMel *Wolbachia* strain, resulting in a measurable reduction in annual dengue cases [22, 23]. Since then, multiple replacement and suppression initiatives have been carried out across Colombia, Brazil, and Vietnam, demonstrating the growing reach of this biocontrol approach [24–26].

Despite these encouraging results, temperature has emerged as a crucial factor affecting the effectiveness of *Wolbachia*-based interventions [27–29]. Experimental work by Ross et al. [28] exposed three *Wolbachia* strains, wMel, wMelPop, and wAlbB, to cyclical temperature conditions ranging from 26°C to 37°C. Under regimes with mean temperatures around 29°C, the densities of wMel and wMelPop within mosquito vectors tended to decline. This is particularly consequential because both cytoplasmic incompatibility and maternal transmission depend on sufficient bacterial density within the host [30, 31]. Of the two mechanisms, maternal transmission appears especially vulnerable, often becoming ineffective or lost entirely following exposure to elevated temperatures. At the same time, cytoplasmic incompatibility diminishes more gradually, a pattern observed across multiple Aedes species [30–32].

The real-world implications of this thermal sensitivity became apparent in 2018, when a heatwave exceeding 40°C for three consecutive days was recorded in Cairns, the very site of the original *Wolbachia* release [33]. This extreme event raised serious concerns about the long-term stability of *Wolbachia* infection frequencies in mosquito populations under increasingly variable climate conditions.

Field successes amid thermal vulnerability have motivated efforts to develop models to predict when *Wolbachia* interventions succeed. Various models of *Ae. aegypti* population dynamics and *Wolbachia* releases have been developed to evaluate ecological and epidemiological impacts [34–37]. Many of these models include temperature-dependent effects, highlighting the crucial role of environmental factors in influencing mosquito and *Wolbachia* behavior [38]. Evaluating intervention effectiveness requires considering the climatic variability of target regions, including seasonal changes and anomalies such as El Niño and extreme heat. There is limited data on how such variability affects *Wolbachia*’s effectiveness and infection dynamics, highlighting a gap between controlled experiments and real-world thermal conditions in dengue-endemic areas.

To address this gap, we develop a mathematical model that integrates temperature-dependent *Wolbachia* parameters across regimes representative of seasonal and interannual climatic variability. We use this framework to examine how temperature patterns and release strategies influence population replacement dynamics, *Wolbachia* persistence, and dengue transmission. We hypothesize that elevated temperatures delay *Wolbachia* establishment and reduce its long-term stability, with downstream consequences for dengue dynamics, given dengue viruses’ dependence on *Wolbachia* frequency to suppress DENV transmission. If confirmed, this would argue for tailoring *Wolbachia* release programs to the local temperature regime, rather than applying a uniform deployment strategy across climatically distinct regions.

## Materials and methods

### Model description

We developed a mathematical model integrating human and vector epidemiological dynamics with temperature-dependent parameters for *Wolbachia*. The mosquito population was divided into two compartments: non-*Wolbachia* (*u*) and *Wolbachia*-infected (*w*), each further split into susceptible and DENV-infected states (*S*_*u*_, *I*_*u*_, *S*_*w*_, *I*_*w*_), assuming a 1:1 female-to-male ratio. The human population (*H*) followed a classical Susceptible–Infectious–Recovered (S, I, R) structure. Fig 1 shows the model schematic.

**Fig 1.**
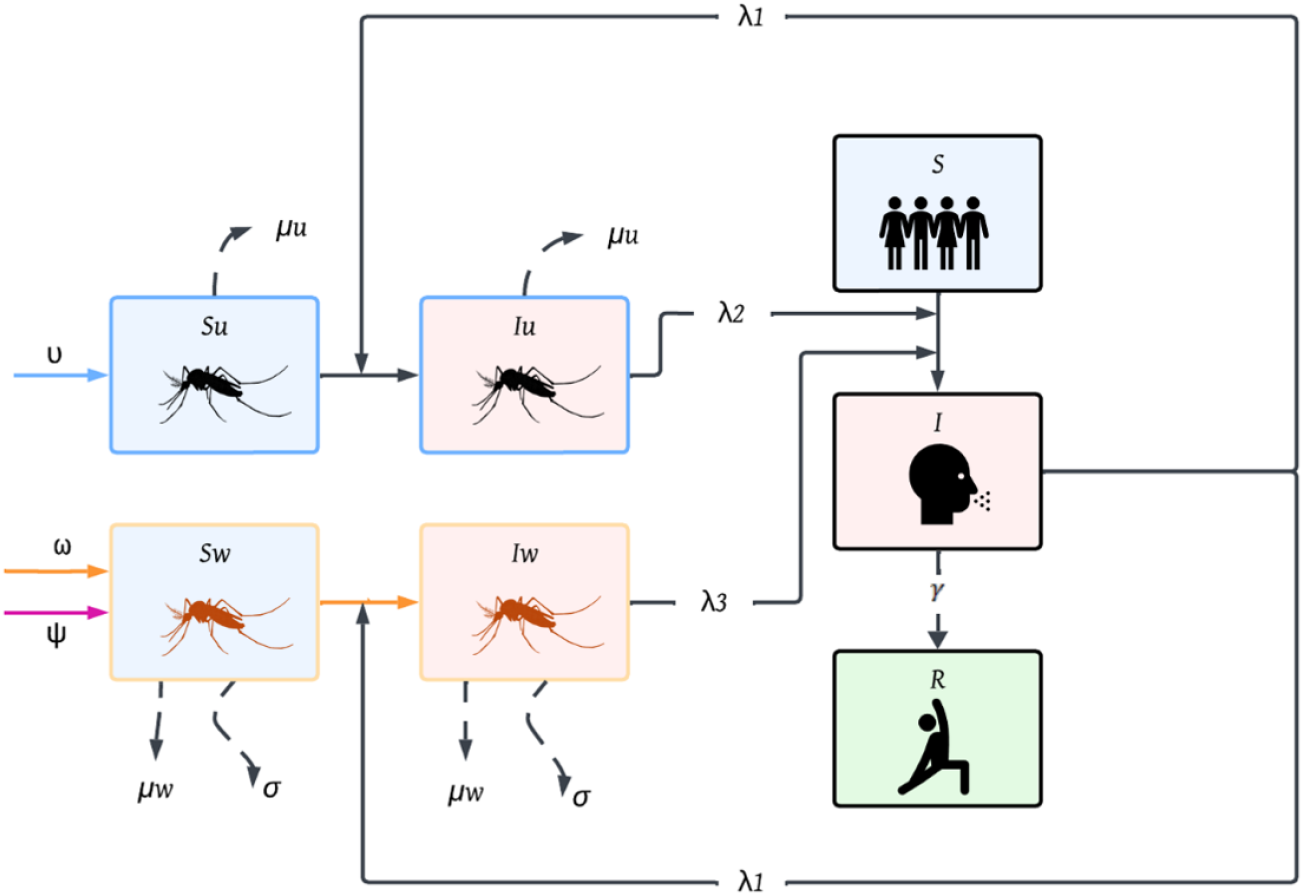
Proposed compartmental model for dengue transmission and *Wolbachia* interventions: The model incorporates two co-circulating mosquito populations: >wild-type uninfected mosquitoes (*S*_*u*_, *I*_*u*_) and *Wolbachia*-infected mosquitoes (*S*_*w*_, *I*_*w*_). Each vector population is structured into susceptible and infectious stages. Human dynamics follow an SIR framework with susceptible (*S*), infectious (*I*), and recovered (*R*) classes. Three force-of-infection terms govern transmission between vectors and humans (*λ*_1_, *λ*_2_, *λ*_3_), capturing contributions from each infectious vector class. Key demographic processes include mosquito recruitment (*v, ω, ϕ*), natural mortality (*µ*_*w*_), *Wolbachia*-induced mortality and reproductive costs (*σ*), and human recovery (*γ*).

The model assumes a constant human population and circulation of a single dengue serotype, which confers complete immunity upon recovery. Non-*Wolbachia* mosquitoes increase at total recruitment rate *υ*, which depends on both infected and uninfected mosquito densities and includes offspring from uninfected-uninfected matings as well as uninfected female–infected male matings under incomplete cytoplasmic incompatibility (*ϕ <* 1). Cytoplasmic incompatibility (CI) is frequency-dependent: higher *Wolbachia* frequencies (*F*_*w*_) increase the likelihood of infected male–uninfected female matings, given sufficient uninfected females, and this is captured through *ϕ*. A small fraction of matings between infected females and either infected or uninfected males also produce uninfected offspring when maternal transmission is incomplete (*ζ <* 1).

*Wolbachia*-infected mosquitoes increase through matings between infected females and males of either type, at total recruitment rate *ω*. The intervention is represented by *ψ*, which varies in magnitude and release interval. Both mosquito populations have distinct mortality rates, *µ*_*u*_ and *µ*_*w*_, with *µ*_*u*_ *< µ*_*w*_. The full system is:

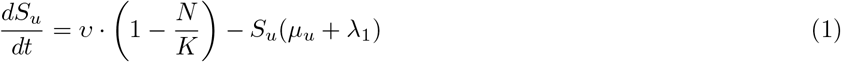

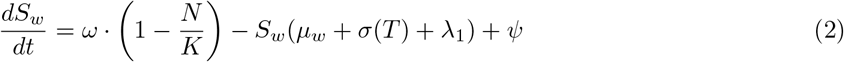

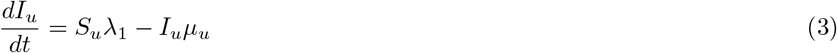

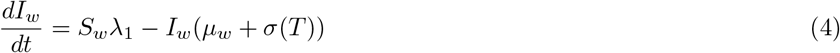

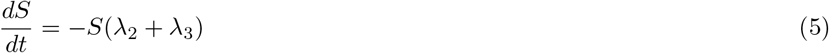

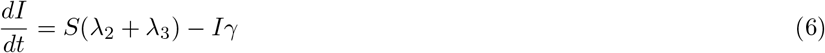

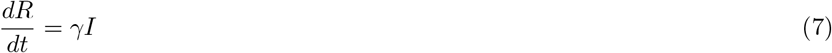

Where

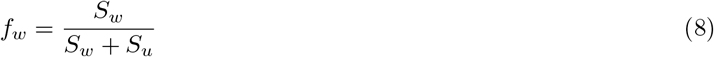

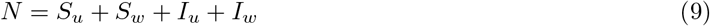

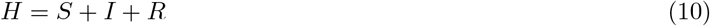

Mosquito recruitment depends on the reproductive rates *r*_*u*_ and *r*_*w*_ of uninfected and *Wolbachia*-infected females, with *r*_*u*_ *> r*_*w*_ reflecting the fitness cost of *Wolbachia* infection [39]. The total recruitment rates *υ* and *ω* incorporate logistic density dependence through carrying capacity *K*, cytoplasmic incompatibility through *ϕ*(*T* ), and maternal transmission through *ζ*(*T* ), the proportion of offspring inheriting the infection:

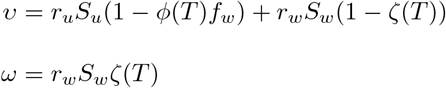

*Wolbachia*-infected mosquitoes also carry a temperature-dependent mortality cost, *σ*(*T* ), reflecting thermal stress associated with infection [40]. DENV transmission between compartments is driven by the *Aedes aegypti* biting rate, *b*_*r*_. Susceptible mosquitoes become infected from infectious humans at rate:

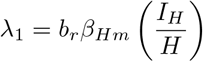

where *β*_*Hm*_ is the human-to-mosquito transmission probability. Susceptible humans face two forces of infection depending on the infection status of the biting mosquito:

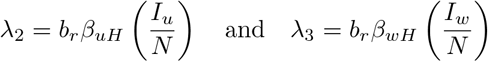

where *β*_*uH*_ and *β*_*wH*_ are the mosquito-to-human transmission probabilities from uninfected and *Wolbachia*-infected mosquitoes, respectively. The lower transmission probability for *Wolbachia*-infected mosquitoes (*β*_*wH*_ *< β*_*uH*_ ) reflects the pathogen-blocking effect central to this biocontrol strategy. Infected humans recover at rate *γ*; *f*_*w*_, *N*, and *H* track *Wolbachia* frequency in the mosquito population, total mosquito abundance, and total human population size, respectively.

### Model parametrization

Field releases used the wMel strain of *Wolbachia pipientis*, which induces moderate cytoplasmic incompatibility in *Aedes aegypti*, is maternally transmitted with high fidelity, and imposes comparatively low fitness costs on the host [41]. The model accounts for wMel’s thermal sensitivity through a temperature-dependent function reflecting reduced CI strength and maternal transmission at higher temperatures. All strain-specific parameters, CI penetrance, maternal transmission efficiency, and effects on adult survival and fecundity, were drawn from laboratory and field studies of wMel (Table 1).

**Table 1.**
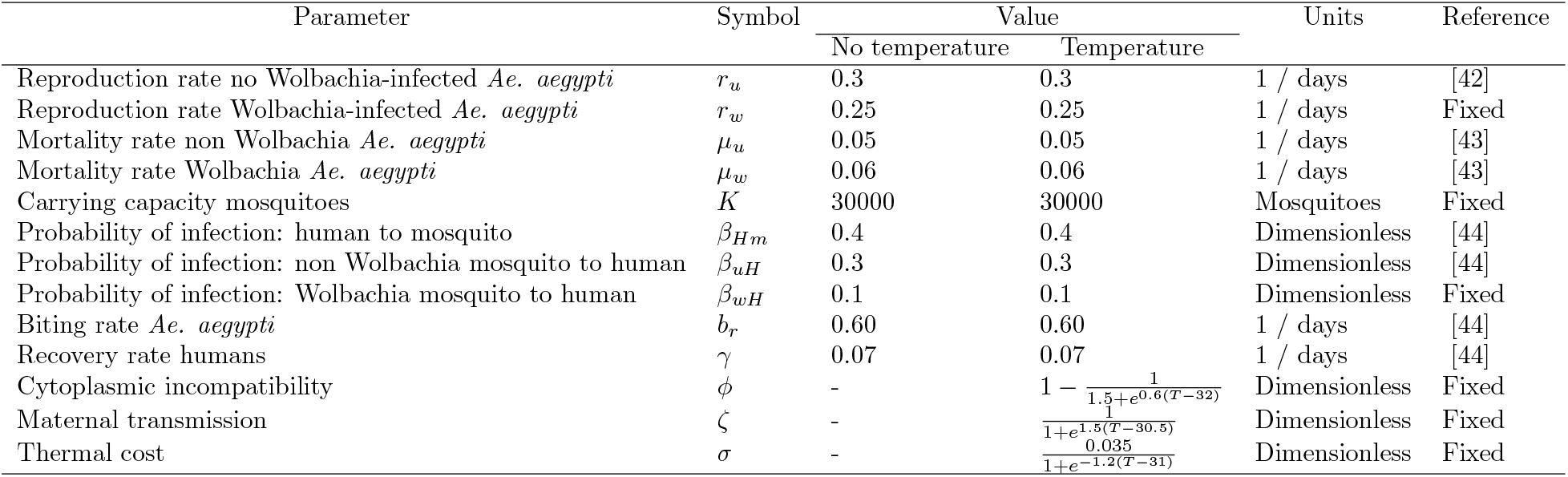
Parameter descriptions, values, and references used in the model.

| Parameter | Symbol | Value |  | Units | Reference |
| --- | --- | --- | --- | --- | --- |
|  |  | No temperature | Temperature |  |  |
| Reproduction rate no Wolbachia-infected *Ae. aegypti* | $r_u$ | 0.3 | 0.3 | 1 / days | [42] |
| Reproduction rate Wolbachia-infected *Ae. aegypti* | $r_w$ | 0.25 | 0.25 | 1 / days | Fixed |
| Mortality rate non Wolbachia *Ae. aegypti* | $\mu_u$ | 0.05 | 0.05 | 1 / days | [43] |
| Mortality rate Wolbachia *Ae. aegypti* | $\mu_w$ | 0.06 | 0.06 | 1 / days | [43] |
| Carrying capacity mosquitoes | $K$ | 30000 | 30000 | Mosquitoes | Fixed |
| Probability of infection: human to mosquito | $\beta_{Hm}$ | 0.4 | 0.4 | Dimensionless | [44] |
| Probability of infection: non Wolbachia mosquito to human | $\beta_{uH}$ | 0.3 | 0.3 | Dimensionless | [44] |
| Probability of infection: Wolbachia mosquito to human | $\beta_{wH}$ | 0.1 | 0.1 | Dimensionless | Fixed |
| Biting rate *Ae. aegypti* | $b_r$ | 0.60 | 0.60 | 1 / days | [44] |
| Recovery rate humans | $\gamma$ | 0.07 | 0.07 | 1 / days | [44] |
| Cytoplasmic incompatibility | $\phi$ | - | $1 - \frac{1}{1.5 + e^{0.6(T-32)}}$ | Dimensionless | Fixed |
| Maternal transmission | $\zeta$ | - | $\frac{1}{1 + e^{1.5(T-30.5)}}$ | Dimensionless | Fixed |
| Thermal cost | $\sigma$ | - | $\frac{0.035}{1 + e^{-1.2(T-31)}}$ | Dimensionless | Fixed |

### Temperature-dependent *Wolbachia* parameters

Cytoplasmic incompatibility (*ϕ*), maternal transmission (*ζ*), and thermal cost (*σ*) are modeled as temperature-dependent functions, parameterized from experimental evidence of reduced *Wolbachia* density in *Aedes aegypti* above 29°C, with accelerated declines near 30°C and 32°C [27–30, 32]. Maternal transmission was assigned a steeper temperature-response curve and a lower inflection point than cytoplasmic incompatibility, consistent with its greater empirical sensitivity to thermal stress [27–30, 32]. *ϕ* and *ζ* are bounded on [0,1] (complete to absent CI and vertical transmission, respectively), and *σ* ranges from 0 to a maximum of 0.035. Functional forms for all three parameters are given in Supplementary Equations S1–S3 (S1 Text), with values across 20–40°C shown in S1 Fig.

### Temperature-driven analyses

Temperature is a key environmental driver of *Wolbachia* establishment, vector population dynamics, and dengue transmission. We evaluated *Wolbachia* invasion dynamics across three thermal scenarios reflecting ecologically relevant patterns of temperature variation: constant temperature, representing environments close to the annual mean with minimal variation; seasonal fluctuation, capturing predictable intra-annual cycles; and interannual variability, representing different year-to-year variation in baseline conditions. Together, these scenarios allow a systematic assessment of how thermal regime shapes long-term *Wolbachia* establishment and persistence under field-like conditions.

#### Seasonal analysis

Seasonality was modeled as a multi-harmonic sinusoidal function with an annual mean of 25°C and amplitude of ±5°C (range 20–30°C), typical of tropical and subtropical environments where *Wolbachia* interventions occur (Eqs S4–S6 in S1 Text; S2 Fig). Three regimes were considered: Regime A, bimodal with a warmer second peak; Regime B, bimodal with a warmer first peak; and Regime C, unimodal with a single heat peak. These regimes differ in the shape and asymmetry of the temperature cycle, allowing evaluation of how thermal structure affects *Wolbachia* establishment and population replacement.

#### Interannual analysis

Long-term climate variation was simulated by modifying the seasonal function to represent three interannual change scenarios (S3 Fig; Eqs S7–S9 in S1 Text): a linear increase in baseline temperature (*λ*), progressive expansion of seasonal amplitude (*α*), and temporal phase displacement (*ω*). Parameter rates, derived from historical trends and scaled for future scenarios [45–47], were used in a sensitivity analysis via bootstrapping across parameter ranges. Mean trajectories and 95% confidence intervals were calculated to assess uncertainty in *Wolbachia* establishment under different environmental pressures.

### Release strategies and simulation scenarios

All simulations began with 25,000 susceptible wild-type mosquitoes and 300 susceptible humans. *Wolbachia* release strategies varied in magnitude (500–3,000 mosquitoes, in increments of 500) and interval (5, 7, 10, and 14 days), yielding a factorial design across thermal scenarios (S1 Table).

Simulations spanned three levels of thermal complexity. Strategies were first tested at four constant temperatures (25, 28, 29, 30°C) over 365 days, using time to reach 60% *Wolbachia* frequency as the primary success measure (replacement threshold), alongside a 20% invasion threshold below which infection decays and the population fails to persist. Each strategy was then simulated under the three seasonal regimes (S2 Fig) to compare replacement dynamics, and a 7-day, 1,500-mosquito release strategy was simulated over two years — releases in year one, none in year two — to evaluate post-release *Wolbachia* persistence (S4 Fig). Finally, strategies were assessed under the three climate change scenarios (S3 Fig) using 100 simulations over five years, focusing on stability of mean *Wolbachia* frequency.

A temperature-dependent SIR model was then used to evaluate dengue infection dynamics under the simulated release scheme. Simulations started from a fully susceptible human population, with three infected individuals introduced on day 90 to simulate viral importation. A fixed intervention of 1,500 *Wolbachia*-infected mosquitoes per week was applied to assess epidemiological outcomes across thermal scenarios, and the resulting dengue peak was compared against a reference scenario under identical conditions without *Wolbachia* releases.

## Results

Based on the proposed temperature-dependent model, we determined that *Wolbachia* establishment and persistence declined as temperature increased across all simulated scenarios, with direct consequences for dengue suppression. Temperature strongly constrained which release strategies achieved 60% population replacement, progressively eliminating viable combinations as conditions warmed (Fig 2). At 25°C (panel A), all strategies with 1,000 or more mosquitoes succeeded, with replacement time decreasing with higher release magnitude and shorter interval. The most intensive strategy (3,000 mosquitoes every 5 days) reached the threshold by day 36; intermediate combinations (1,000–2,500 mosquitoes at 7-day intervals) required 57–132 days. Releases of 500 mosquitoes failed at all intervals at this temperature. At 28°C (panel B), the same pattern held, but replacement times increased across all viable strategies, with the strongest intervention requiring approximately 41 days and the 1,000-mosquito strategies extending to 64–159 days. The 500-mosquito releases continued to fail, and longer intervals ( ≥10 days) at 1,000 mosquitoes also dropped below the replacement threshold. At 29°C (panel C), failure expanded further: all 500-mosquito releases and most low-to intermediate-magnitude strategies at intervals longer than 7 days failed to reach the target. Only high-magnitude releases ( ≥2,000 mosquitoes at 5-day intervals, or 3,000 mosquitoes at up to 10-day intervals) remained effective, with the strongest strategy crossing the threshold by day 46. At 30°C (panel D), only the single most intensive combination, 3,000 mosquitoes released every 5 days, achieved replacement. All other strategy combinations failed, reflecting the combined suppression of *Wolbachia* maternal transmission and cytoplasmic incompatibility at this temperature.

**Fig 2.**
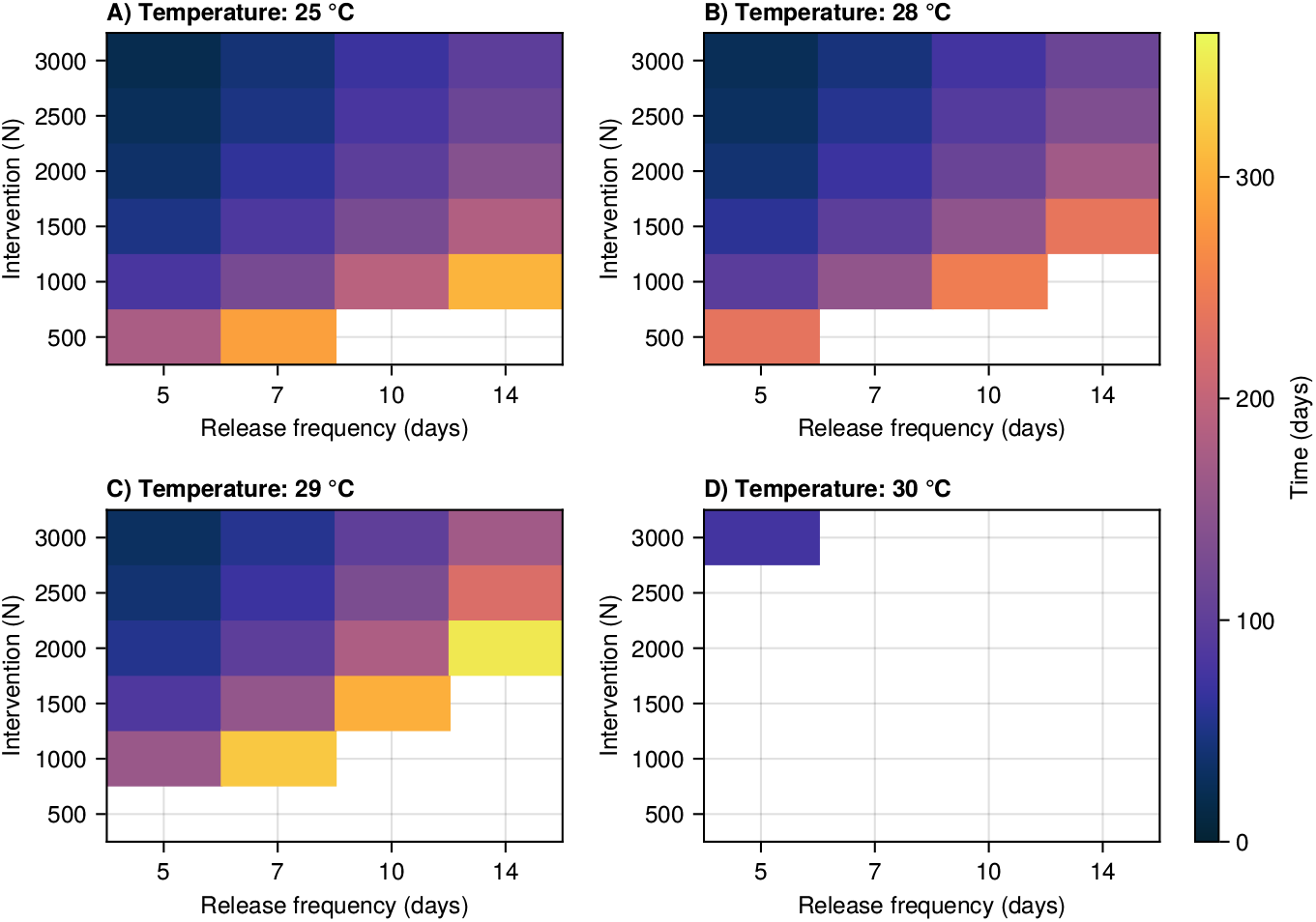
Replacement times under different intervention strategies simulated at constant temperatures (25 °C, 28 °C, 29 °C, and 30 °C). Each strategy combines a release magnitude and a release interval. White cells indicate scenarios that did not reach the Wolbachia replacement threshold (60%)

Seasonal temperature profiles influenced both the speed and success of population replacement across all three regimes (Fig 3). Regime C was the most lenient, allowing replacement across nearly the entire parameter space, including strategies like 500-mosquito releases every 5 days, and achieving the fastest replacement times among them. A noticeable diagonal gradient in this regime indicates the combined effect of release magnitude and interval, with replacement times increasing from less than 50 days at high magnitude and short intervals to over 200 days at mid-range combinations. Regime A was moderate: high-magnitude approaches replaced populations in about 100 days, but the 500-mosquito strategy failed at intervals of 7 days or more. Regime B was the most restrictive, requiring the longest replacement times overall, with intermediate strategies (1000–2500 mosquitoes released every 7–14 days) taking an average of 171 days to achieve successful replacement, delaying *Wolbachia* establishment by nearly 47 days relative to Regime A.

**Fig 3.**
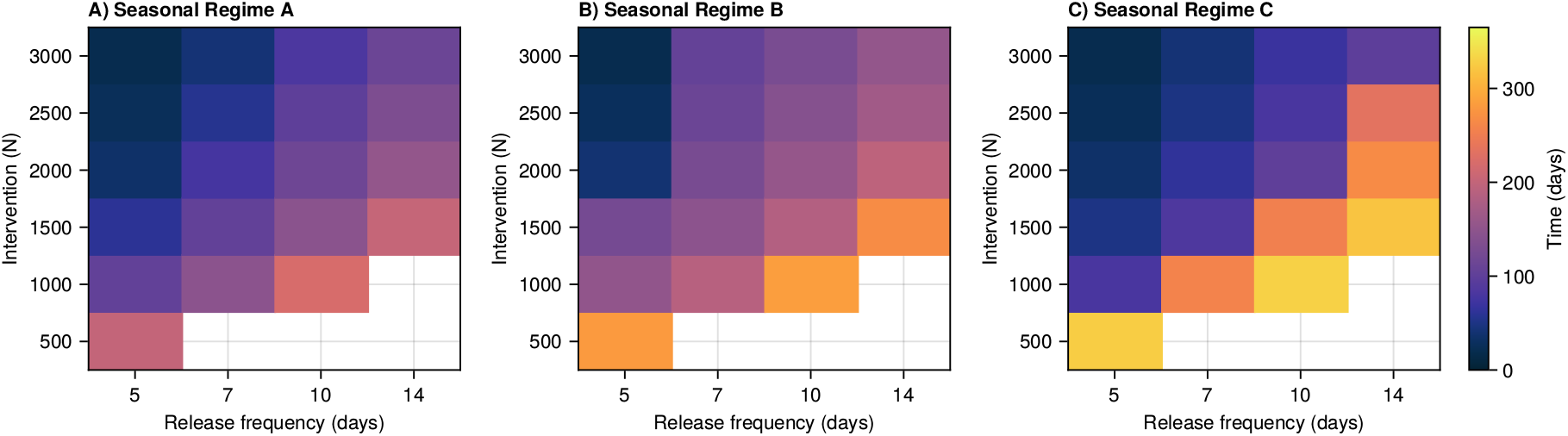
Replacement times under the three seasonal temperature regimes (Regime A, B, and C). Each strategy combines a release magnitude and a release interval. White cells indicate scenarios that did not reach the *Wolbachia* replacement threshold (60%).

*Wolbachia* frequency showed consistent seasonal fluctuations across all three interannual temperature scenarios. Still, the type of thermal disturbance led to notably different long-term patterns (Fig 4). Because the interannual scenarios were bootstrapped over 100 runs, the shaded areas represent the 95% confidence intervals around the mean *Wolbachia* frequency trajectories. In the case of a progressive baseline warming trend (panel A – baseline change), the oscillation peaks decreased from about 1.0 in year 1 to around 0.8 by year 5. At the same time, the troughs dropped from approximately 0.35 to 0.15, nearing the 20% invasion threshold by the end of the period. This scenario also demonstrated the widest confidence intervals, indicating high sensitivity to thermal fluctuations. With seasonal intensity expansion (panel B – intensity change), peak frequencies reliably returned to roughly 1.0 after each thermal stress, and troughs stabilized near 0.2, close but not below the invasion threshold, showing no overall trend over five years and maintaining narrow confidence intervals. Phase displacement (panel C – phase displacement change) also produced a stable oscillatory pattern, with peaks at about 1.0 and troughs between 0.2 and 0.3 but with a distinctly asymmetric waveform, frequencies increased sharply from troughs and declined rapidly from peaks, contrasting with the smoother oscillations seen in the baseline and in the intensity change (Fig. 4A and 4B). Confidence intervals for the phase displacement change (Fig. 4C) were the narrowest among the three scenarios.

**Fig 4.**
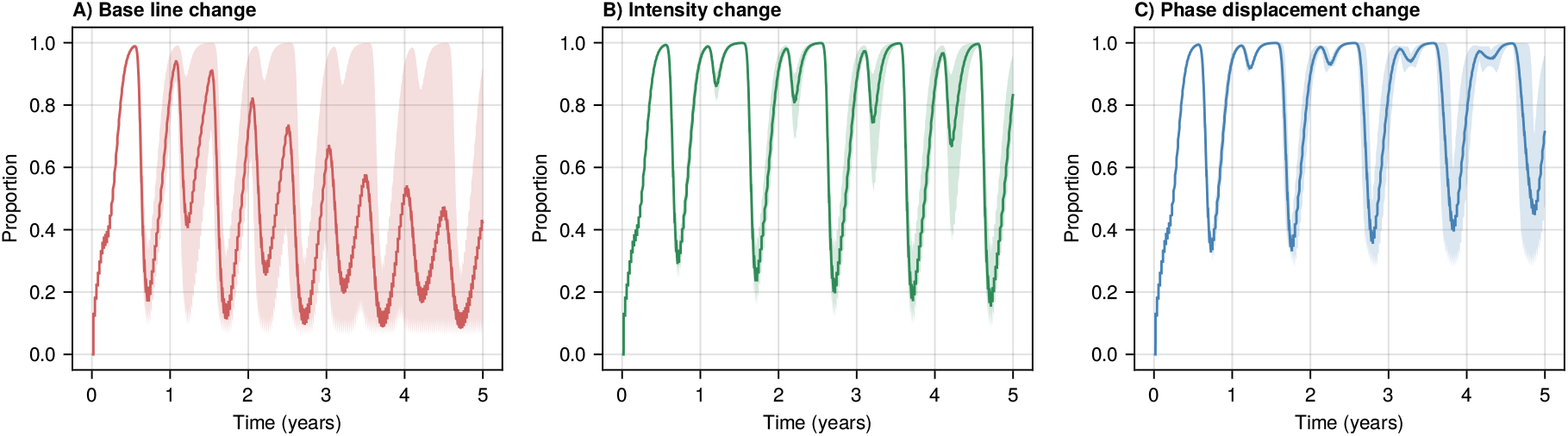
*Wolbachia* frequency under interannual temperature trajectories. Representative intervention: 1,500 mosquitoes released every 7 days, within a one-year simulation. (A) Baseline scenario (Eq S7), (B) Intensity change scenario (Eq S8), (C) Phase displacement scenario (Eq S9).

Epidemiological SIR simulations combined with *Wolbachia* dynamics showed that outbreak severity increased with higher temperatures due to reduced *Wolbachia* establishment. At the same time, peak infection occurred earlier as temperatures rose (Fig 5). The reference scenario, without *Wolbachia* intervention, produced a dengue peak of 107 cases on day 132, approximately 36% of the simulated population. At 25°C (SIR - panel A), *Wolbachia* frequency reached nearly 1.0 around day 150, resulting in a dengue peak of about 48 infections on day 161 (16% of the simulated population), which corresponds to a 54% reduction compared to the reference scenario. At 28°C (SIR-panel B), *Wolbachia* established more slowly, reaching nearly 1.0 by day 175; meanwhile, the dengue peak rose to about 60 cases at day 151, representing 20% of the total population, and a lower infection reduction (44%) relative to the reference scenario. This indicates a longer period of incomplete *Wolbachia* coverage during virus transmission. At 29°C (SIR - panel C), *Wolbachia* plateaued at roughly 85% without fully replacing the population, resulting in a dengue peak of about 81 cases around day 150 (27% of the simulated population), corresponding to an estimated 24% reduction relative to the reference scenario. At 30°C (SIR - panel D), *Wolbachia* did not establish at all, instead oscillating around 20–25%, declining gradually without reaching zero within the simulation; the earliest and largest dengue peak occurred around day 136 with 96 cases, (32% of the simulated population), representing the minimal infection reduction of 10%. In all cases, peak infection occurred earlier as temperature increased, from day 161 at 25°C to day 136 at 30°C, corresponding to the reduced *Wolbachia* frequency at virus introduction.

**Fig 5.**
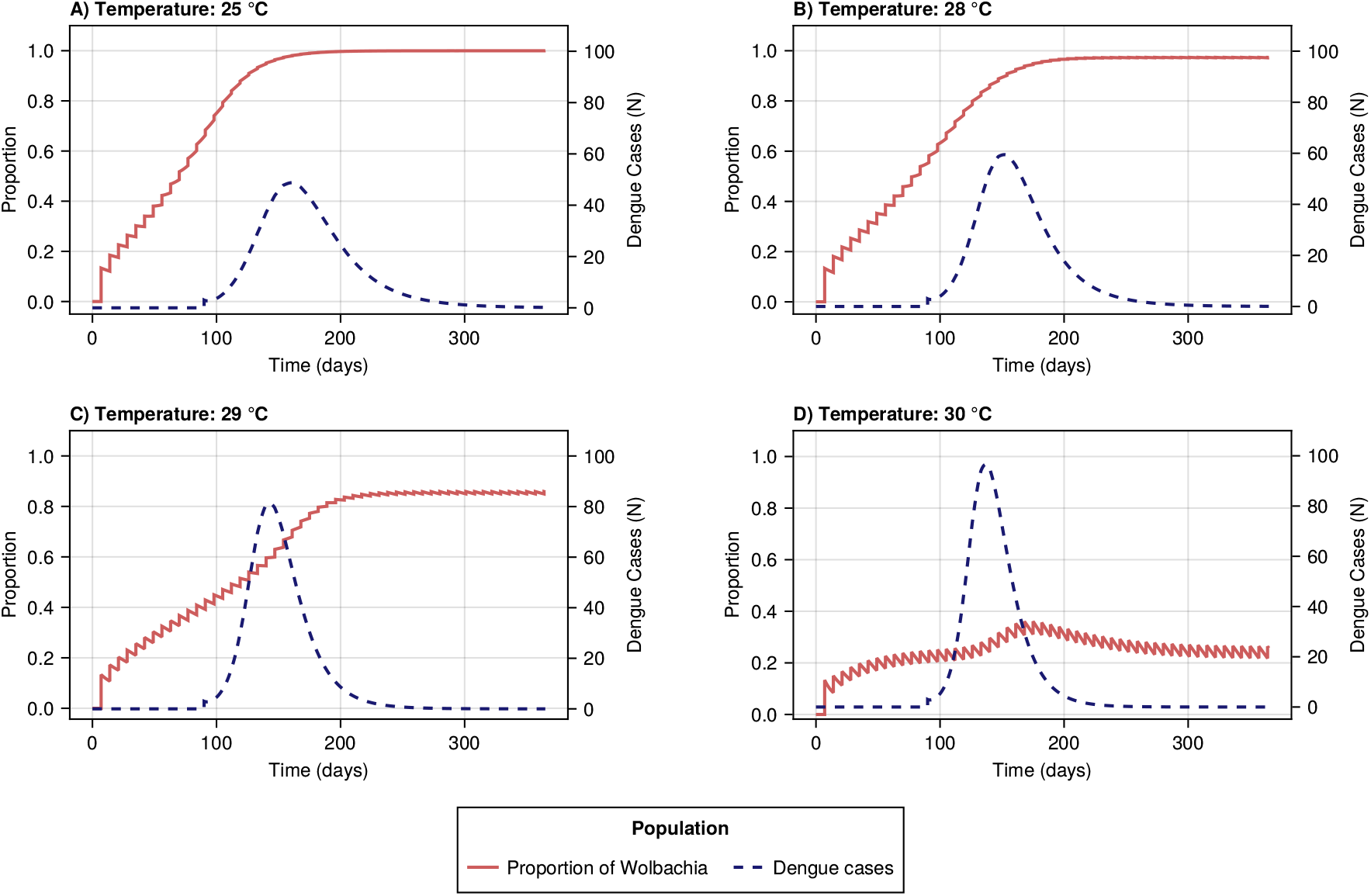
Dynamics of *Wolbachia* frequency and infected humans under constant temperature scenarios (25 °C, 28 °C, 29 °C, and 30 °C). The left axis represents *Wolbachia* frequency (red solid line), and the right axis represents the number of infected humans (blue dashed line). DENV introduction on day 90 with constant human population of N = 300.

## Discussion

This study evaluated how temperature shapes the establishment, persistence, and epidemiological impact of *Wolbachia*-based vector control strategies against dengue. At constant temperatures, the reduction in viable strategies between 28°C and 30°C reflects the well-documented decrease in *Wolbachia* density within *Ae. aegypti* at elevated temperatures, particularly for the wMel strain, which impairs both maternal transmission and cytoplasmic incompatibility and thereby weakens the reproductive advantage of infected mosquitoes [28, 29, 31, 32]. In our model, maternal transmission declined more steeply than cytoplasmic incompatibility as temperature rose from 28°C to 30°C (24% versus 14% reduction, respectively), consistent with experimental evidence of differential thermal sensitivity between these two mechanisms [27–30, 32]. Our simulations produced replacement times of 71-105 days at 25-28°C with comparable field intensities, and only the most intensive strategy (3,000 mosquitoes every five days, approximately 70,000 mosquitoes/km^2^ per week) succeeded at 30°C, placing it at the upper bound of typical field deployments (20,000-60,000 mosquitoes/km^2^ per week [23–26]). Lopes et al. [35] similarly showed that *Wolbachia* prevalence declined linearly with rising temperature in modeled mosquito populations, and that above approximately 28.9°C, intervention efficacy fell below 50% regardless of release intensity. For program design in warm climates, this is consequential: both maternal transmission and cytoplasmic incompatibility operate under continuous thermal stress with no seasonal window for recovery, demanding larger releases and shorter intervals than cooler settings to offset the constant fitness disadvantage relative to wild-type mosquitoes. Seasonality compounds this picture beyond mean temperature or amplitude alone: our simulations produced an oscillatory dynamic in which *Wolbachia* frequency fell during thermal peaks and rebounded during cooler periods. Releases launched too close to the period of maximum thermal stress met *Wolbachia* before it reached a protective frequency, requiring more releases and a longer path to population replacement; releases launched well ahead of the hottest period instead let *Wolbachia* climb to higher frequencies and recover more readily afterward—a pattern mirrored in Vietnam, where establishment succeeded at cooler sites but failed to stabilize at hotter ones [26]. Initiating releases too close to the seasonal peak therefore raises both intervention cost and the risk of outright failure, underscoring that deployment timing must treat thermal conditions as a dynamic constraint, not merely a static backdrop.

Regarding interannual climate variability, the results highlight the importance of extrapolating these interventions to future climate conditions. The progressive increases in baseline temperature generated wider uncertainty intervals in *Wolbachia* frequency, reflecting the high sensitivity to the rate of mean warming. As mean temperature rises, the parameters of cytoplasmic incompatibility and maternal transmission approach their minimum values progressively, limiting *Wolbachia*’s ability to return to higher frequencies between oscillatory cycles. Despite a clear downward trend over the simulated period, infection was not completely lost across any simulation. Under seasonal intensity expansion, *Wolbachia* exhibited stable oscillatory dynamics, with sharper, more regular declines as the seasonal intensity widened. This is consistent with observations from Cairns, Australia, where a heatwave reaching approximately 43°C caused a reduction in wMel frequencies that subsequently recovered within months [33], and where the duration and timing of thermal stress, rather than its intensity, appeared to be the primary determinants of population stability. Temporal phase displacement generated a misalignment between *Wolbachia* frequency and thermal stress. Because the timing of the thermal peaks shifts progressively, heat stress coincides with different phases of the *Wolbachia* frequency cycle. Although phase displacement alone did not prevent *Wolbachia* frequency recovery under the perturbation magnitudes tested, cumulative phase displacement shortens the thermal recovery window between consecutive heat peaks, exposing *Wolbachia* populations to successive stress events before frequency can rebound. Critically, a population can decline below the invasion threshold by a heat peak with unchanged intensity if that peak arrives earlier in the annual cycle. Taken together, these results suggest that release programs must monitor and detect declining frequency trends before populations fall below the invasion threshold, and deployment programs in regions projected to experience significant warming should evaluate more intense deployments or alternative strains with greater thermal tolerance.

Temperature also shaped dengue outbreak dynamics under the simulated intervention. At 25°C and 28°C, *Wolbachia* frequency was high at the time of virus introduction, limiting the proportion of competent vectors and effectively suppressing transmission, with the greatest dengue reduction observed at 25°C (54%). At 29°C and 30°C, *Wolbachia* frequency at the time of virus introduction was approximately 38% and 20%, respectively, resulting in larger outbreaks that peaked earlier. At 30°C, the reduction in dengue transmission fell to 10%, reflecting the combined effect of incomplete population replacement and the temperature-dependent reduction in *Wolbachia*’s ability to suppress DENV dissemination within the vector [48, 49]. Even more, the introduction of a virus during a thermal peak, when *Wolbachia* frequency is at its lowest, will produce a substantially larger outbreak than one introduced during a cooler period or under stable conditions, such as the 25 °C scenario. These findings are consistent with field reductions reported in Brazil, Colombia, Indonesia, and Australia [7, 23, 26, 50]. In Rio de Janeiro, *Wolbachia* frequencies above 60% were associated with a 76% reduction in dengue cases [26, 50], and in Medellín, Colombia, where the annual temperature profile is nearly isothermal at approximately 22–23°C, a 95% reduction in incidence was reported [51], illustrating how thermally stable environments can sustain both high *Wolbachia* frequencies and effective dengue suppression. It should be noted that there are uncertainties regarding the epidemiological impact of *Wolbachia* interventions, as field evidence suggests that observed reductions in dengue incidence may be related to broader factors rather than the intervention itself [52]. In Niterói, Brazil, where a 69–90% reduction in dengue incidence was reported, the decline was partially attributable to vaccination campaigns and vector control measures, and similar reductions were observed in other cities in the state of Rio that did not receive *Wolbachia* intervention during the same period [52]. These observations reflect the complexity of attributing epidemiological outcomes solely to *Wolbachia* deployments in regions where multiple control measures operate concurrently. Furthermore, the effect of temperature on DENV blockage within *Ae. aegypti* remains partially known, which limits the epidemiological inferences that can be made from our simulations; rising temperatures may reduce this blockage, allowing *Wolbachia*-infected mosquitoes to transmit dengue at higher rates than current models predict.

Several modeling assumptions warrant consideration when interpreting these results. The omission of aquatic life stages (eggs, larvae, and pupae) represents the most consequential structural limitation, as these stages are highly sensitive to environmental conditions and influence recruitment and population dynamics in ways not captured here. Mosquito life-history parameters were also treated as fixed rather than temperature-dependent, which may overestimate the contribution of *Wolbachia*-specific thermal effects by holding background mosquito biology constant. Incorporating temperature-dependent life-history parameters would improve model realism. The temperature-dependent functions for maternal transmission, cytoplasmic incompatibility, and thermal cost were constructed from laboratory experiments conducted under controlled conditions and may not fully reflect field behavior. Additionally, the assumption that DENV-infected mosquitoes do not reproduce may underestimate population growth, and the assumption of complete immunity following dengue infection oversimplifies multi-serotype dynamics. Despite these limitations, the results carry practical implications for intervention planning. The timing of releases relative to the seasonal temperature profile is a critical operational consideration: initiating releases during or immediately before peak temperatures may slow establishment and reduce the probability of successful population replacement. Once *Wolbachia* reaches high frequencies, simulated populations showed resilience to short-term thermal anomalies, suggesting that long-term stability is achievable under the variable climate conditions typical of tropical deployment settings.

## Conclusion

This study demonstrates that temperature is a decisive factor in *Wolbachia* based interventions, dictating how intensively *Wolbachia* must be deployed to establish itself in the population. In tropical regions with limited seasonal thermal variation, mean temperature alone dominates *Wolbachia* fitness, impairing both cytoplasmic incompatibility and maternal transmission. Seasonality shapes population dynamics, suppressing *Wolbachia* prevalence as temperatures rise, then allowing recovery as they decline. Accordingly, *Wolbachia* deployments should optimally begin before heat peaks, allowing sufficient time for *Wolbachia* establishment and stability before thermal stress intensifies. Critically, the timing of thermal stress relative to population recovery is a key determinant for long-term persistence: when the recovery interval between successive heat periods is insufficient to retrieve high infection frequencies, thermal stress accumulates progressively, lowering *Wolbachia* prevalence and increasing the risk of losing infection across the population. Beyond temperature, additional factors such as vector control, rainfall, and geographical heterogeneity should be integrated into deployment planning, as similar climate conditions can yield markedly different population dynamics.

*Wolbachia*-based interventions nonetheless proved effective in reducing dengue transmission across a broad range of thermal conditions. The release of wMel-infected *Aedes aegypti* mosquitoes substantially suppresses the magnitude of dengue outbreaks, with the greatest reduction observed at 25°C (54.6%). Moreover, higher *Wolbachia* population frequencies delayed the timing of dengue virus peaks, extending the epidemiological response window for public health authorities. However, as temperatures approached the *Wolbachia* persistence threshold ( ∼30°C), outbreaks were both larger and earlier, indicating growing challenges in sustaining intervention efficacy amid rising global temperatures and the increasing frequency of extreme heat events.

## Supporting information

### S1 Text. Temperature-dependent parameters, release strategies, and temperature regimes

Derivation and functional forms of the temperature-dependent *Wolbachia* parameters, the release-strategy design, and the equations underlying the seasonal and interannual temperature regimes used in the simulations.

#### Temperature-dependent parameters

Cytoplasmic incompatibility (*ϕ*) is modeled as a sigmoidal decreasing function of temperature ranging from 0 to 1, where 0 indicates the absence of cytoplasmic incompatibility (i.e., matings between uninfected females and infected males produce viable offspring) and 1 indicates complete cytoplasmic incompatibility:

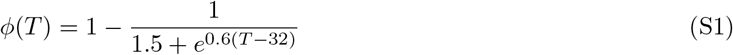

Maternal transmission (*ζ*) is modeled as a sigmoidal decreasing function of temperature, ranging from 0 to 1, where 0 indicates the absence of vertical transmission of *Wolbachia* and 1 represents complete maternal transmission:

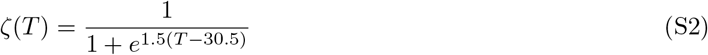

Thermal cost (*σ*) is modeled as an increasing sigmoidal function of temperature, ranging from 0 to 0.035, where *σ* = 0 indicates no thermal cost in adult stages associated with high temperature and *Wolbachia* infection. The maximum value was set to 0.035, corresponding to an approximate 40% reduction in vector performance at high temperatures:

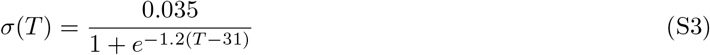

#### Release strategies

Release intervals were selected based on field deployment protocols typically ranging from 5 to 7 days [19, 20, 23]; intervals of 10 and 14 days were included to evaluate less frequent, operationally less demanding schedules. Release magnitudes were determined assuming an urban population density of 5,000 inhabitants/km^2^, under which 300 humans correspond to a simulated area of approximately 0.06 km^2^. The range of magnitudes is consistent with intensities reported in operational *Wolbachia* programmes [19,23,42,43]. The resulting release magnitudes and intervals are summarized in S1 Table.

#### Seasonal temperature regimes

The three seasonal regimes shown in S2 Fig were generated by adjusting the phase and amplitude parameters to produce the bimodal and unimodal profiles described in the main text.

Regime A:

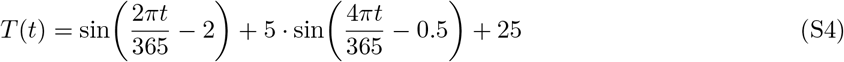

Regime B:

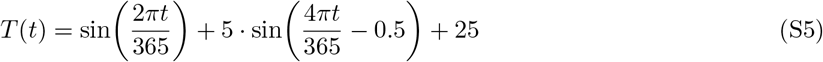

Regime C:

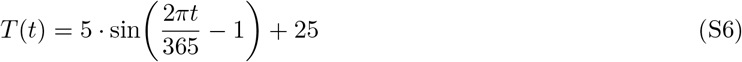

#### Interannual temperature trajectories

Three interannual change scenarios, shown in S3 Fig, were generated by modifying the seasonal function above.

Baseline change:

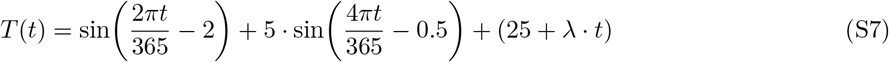

Intensity change:

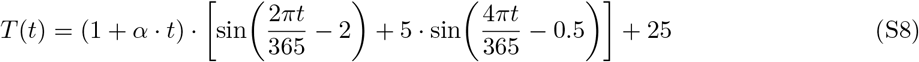

Phase displacement change:

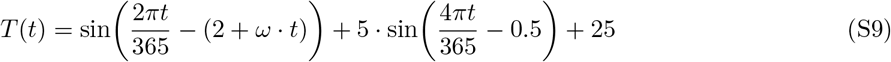

where *λ, α*, and *ω* are rates of change derived from historical climate records and scaled to represent intensified future scenarios [45–47].

**S1 Fig. Temperature-dependent parameters**. Behavior of cytoplasmic incompatibility (*ϕ*), maternal transmission (*ζ*), and thermal cost (*σ*) across the range 20–40°C. Modeled by Eqs S1–S3.

**S2 Fig. Seasonal temperature variability regimes**. Three seasonal scenarios modeled by Eqs S4–S6: Regime A; (II) Regime B; and (III) Regime C.

**S3 Fig. Representative interannual temperature trajectories**. Three interannual scenarios modeled by Eqs S7–S9: (I) Baseline change; (II) Intensity change; and (III) Phase displacement.

**S4 Fig. *Wolbachia* frequency over a two-year period under the three seasonal temperature regimes**. Intervention was simulated with a seven-day release interval, and each colored line represents a different intervention magnitude. Dashed line indicates 60% *Wolbachia* frequency. Dotted line indicates the end of the intervention period.

**S1 Table.**
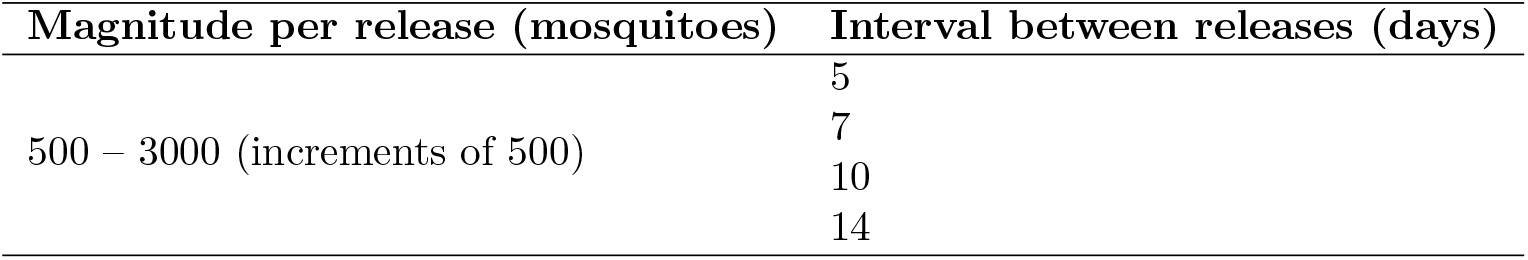
Release magnitudes and intervals implemented in the simulations. Each release magnitude was simulated with all four intervention intervals, yielding a total of 24 release strategies.

## Acknowledgments

This section is intended only for general acknowledgements and thanks. Any information related to funding, data availability, author contributions, etc. should be entered directly into their dedicated fields in the PLOS Editorial Manager submission system, which will then be incorporated into the appropriate section in your article during the production process.

