## Supplementary material for "Temperature variation as a driver of *Wolbachia* release efficacy: implications for dengue control in warming climates": SI

### Supplementary information

#### Temperature-dependent parameters

Temperature-dependent parameters were defined based on experimental evidence, indicating that Wolbachia density decreases within the host at temperatures above 29°C, with accelerate declines at approximately 30°C and 32°C [27-31]. This reduction impairs cytoplasmic incompatibility and maternal transmission, reducing its effectiveness. Additionally, a thermal cost was included to capture the thermal stress associated with carrying the infection at elevated temperatures [40].

Cytoplasmic incompatibility (ϕ) is modeled as a sigmoidal decreasing function of temperature ranging from 0 to 1, where 0 indicates the absence of cytoplasmic incompatibility (i.e., matings between uninfected females and infected males produce viable offspring), and 1 indicates complete cytoplasmic incompatibility.

$$\phi(\mathrm{T}) = 1 - \frac{1}{1.5 + e^{0.6(\mathrm{T}-32)}}) \qquad \text{(Eq S.1)}$$

Maternal transmission (ζ) is modeled as a sigmoidal decreasing function of temperature, ranging from 0 to 1, where 0 indicates the absence of vertical transmission of *Wolbachia* and 1 represents complete maternal transmission.

$$\zeta(T) = \frac{1}{1 + e^{1.5(T-30.5)}} \qquad \text{(Eq S.2)}$$

Thermal cost is modeled as an increasing sigmoidal function of temperature, ranging from 0 and 0.035, where $\sigma = 0$ indicates no thermal cost in adult stages associated with high temperatures and Wolbachia infection. The maximum value was set to 0.035, corresponding to an approximate reduction of 40% in vector performance at high temperatures,

$$\sigma(\mathrm{T}) = \frac{0.035}{1 + e^{-1.2(\mathrm{T}-31)}} \qquad \text{(Eq S.3)}$$

#### Release strategies

Release intervals were selected based on field deployment protocols typically ranging from 5 to 7 days [19, 20, 23]; intervals of 10 and 14 days were included to evaluate less frequent, operationally less intensive schedules. Release magnitudes were determined assuming an urban population density of 5,000 inhabitants/km², under which 300 humans correspond to a simulated area of approximately 0.06 km². The range of magnitudes is consistent with intensities reported in operational Wolbachia programs [19, 23, 42, 43].

**S1 Table. Release magnitudes and intervals implemented in the simulations.** Each release magnitude was simulated with all four intervention intervals, obtaining a total of 24 release strategies.

| Magnitude per release (mosquitoes) | Interval between release (days) |
|---|---|
| 500 – 3000 (increments of 500) | 5 |
| | 7 |
| | 10 |
| | 14 |

#### Seasonal temperature regimes

The three seasonal regimes shown in Fig. S2 were generated using sinusoidal functions with a period of 365 days. Seasonal profiles were obtained by adjusting the phase and amplitude parameters while maintaining the same mean annual temperature (25 °C) and combining annual and semiannual harmonics. Regimes A and B represent bimodal seasonal patterns, with differences in the relative intensity of the two annual heat peaks produced by differences in the phase shift of the annual harmonic. Regime C represents a unimodal seasonal profile generated using only the annual harmonic with a larger amplitude.

##### Regime A:

$$\sin\left(2\pi\frac{t}{365}-2\right)+5\ \cdot\sin\left(4\pi\frac{t}{365}-0.5\right)+25 \qquad \text{(Eq S.4)}$$

##### Regime B:

$$\sin\left(2\pi\frac{t}{365}\right)+5\ \cdot\sin\left(4\pi\frac{t}{365}-0.5\right)+25 \qquad \text{(Eq S.5)}$$

**Regime C:**

$$5 \cdot \sin\left(2\pi \frac{t}{365} - 1\right) + 25 \qquad \text{(Eq S.6)}$$

### Interannual temperature trajectories

To simulate long-term climate variation, three interannual change scenarios were considered, with parameters rates derived from historical climate records and scaled to represent intensified future climate scenarios [45-47]. These interannual trajectories were modeled by progressively modifying the baseline seasonal regime (Regime A): (I) baseline change, in which the mean annual temperature increases at a rate determined by the parameter λ, (II) seasonal intensity change, in which the amplitude of seasonal temperature fluctuations increases progressively at a rate determined by the parameter α, and (III) phase displacement change, in which the timing of the annual harmonic shifts over time at a rate determined by the parameter ω.

**Baseline change:**

$$T(t) = \sin\left(2\pi \cdot \frac{t}{365} - 2\right) + 5 \cdot \sin\left(4\pi \cdot \frac{t}{365} - 0.5\right) + (25 + \lambda \cdot t) \qquad \text{(Eq S.7)}$$

**Intensity change:**

$$T(t) = (1 +\cdot\, \alpha \cdot t) \cdot [\sin\left(2\pi \cdot \frac{t}{365} - 2\right) + 5 \cdot \sin\left(4\pi \cdot \frac{t}{365} - 0.5\right)] + 25 \qquad \text{(Eq S.8)}$$

**Phase displacement change:**

$$T(t) = \sin\left(2\pi \cdot \frac{t}{365} - (2 + \omega \cdot t)\right) + 5 \cdot \sin\left(4\pi \cdot \frac{t}{365} - 0.5\right) + 25 \qquad \text{(Eq S.9)}$$

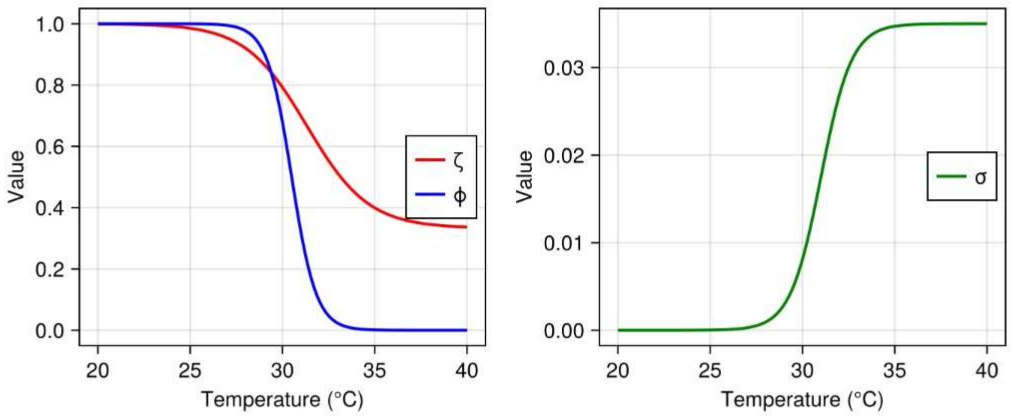


**S1 Fig. Temperature-dependent parameters**
Behavior of cytoplasmic incompatibility (φ), maternal transmission (ζ), and thermal cost (σ) across the range 20–40°C. Modeled by Eqs. S1-S3.

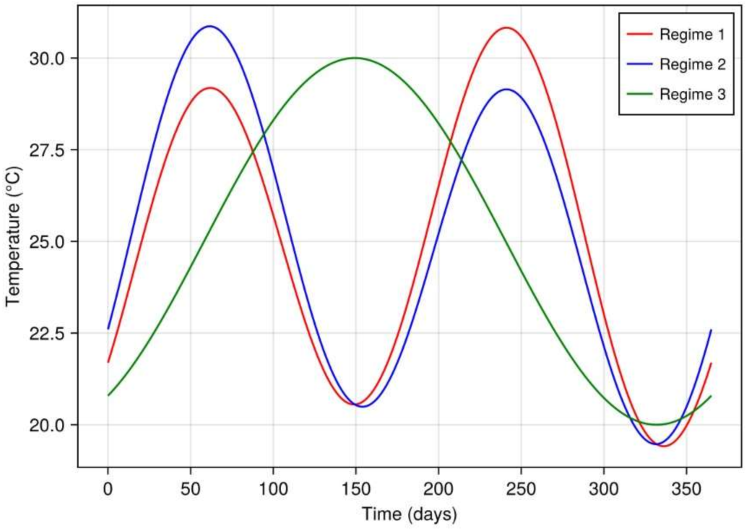


**S2 Fig. Seasonal temperature variability regimes**
Three seasonal scenarios modeled by Eqs. S4-S6: (I) Regime A; (II) Regime B; and (III) Regime C

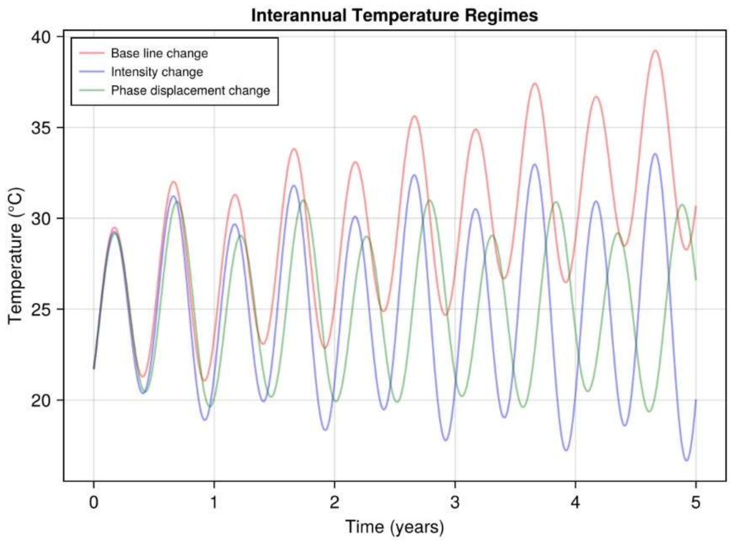


**S3 Fig. Representative interannual temperature trajectories**
Three interannual scenarios modeled by Eqs. S7-S9: (I) Baseline change; (II) Intensity change; and (III) Phase displacement.

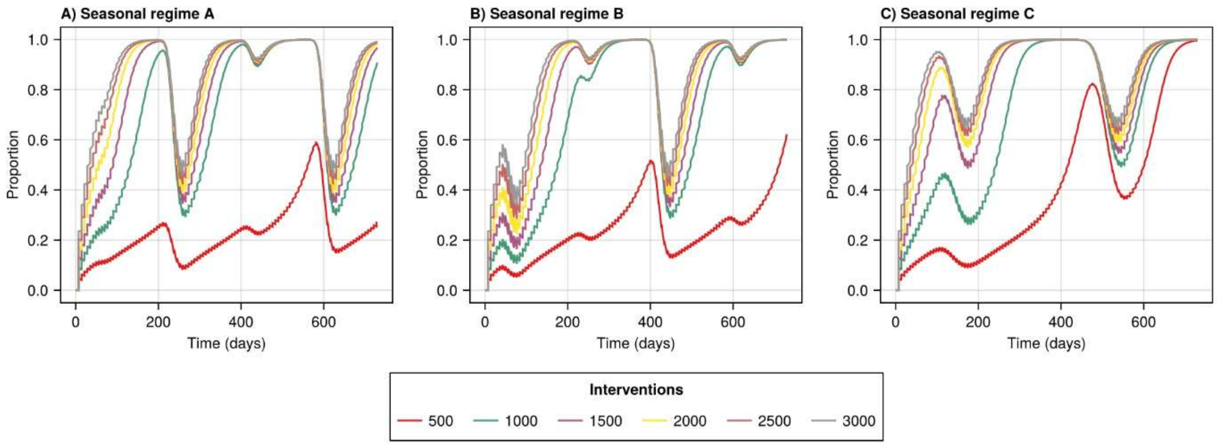


**S4 Fig. Wolbachia frequency over a two-year period under the three seasonal temperature regimes**. Intervention was simulated with a seven-day release interval, and each colored line represents a different intervention magnitude. Dashed line indicates 60% *Wolbachia* frequency. Dotted line indicates the end of the intervention period.
